# Synergistic interactions of mobile genetic elements shape the stepwise evolution of multidrug-resistant plasmids

**DOI:** 10.1101/2025.11.28.691237

**Authors:** Yuqing Mao, Guorui Zhang, Zhiqian Han, Joanna L. Shisler, Rachel J. Whitaker, Thanh H. Nguyen

**Affiliations:** The Grainger College of Engineering, Department of Civil and Environmental Engineering, University of Illinois Urbana-Champaign, IL, USA; Carl R. Woese Institute for Genomic Biology, University of Illinois Urbana-Champaign, IL, USA; Department of Microbiology, University of Illinois Urbana-Champaign, IL, USA; Carle Illinois College of Medicine, University of Illinois Urbana-Champaign, IL, USA

**Keywords:** Integron, IS26, ISCR, antimicrobial resistance, plasmid, mobile genetic element, multidrug resistance

## Abstract

Rising multidrug-resistant (MDR) bacteria threatens global public health. While antibiotic selection preserves MDR plasmids, how mobile genetic elements (MGEs) drive the antibiotic resistance gene (ARG) accumulation to form complex plasmid architectures remain elusive. Here, we demonstrate that MDR plasmid evolution follows a hierarchical scheme, where progressive combinations of integrons, IS26, and ISCR elements drive stepwise increases in ARG load. The combination of all these three MGEs were significantly linked to the highest ARG load on plasmids. In the presence of integrons or ISCR elements, ARG numbers in plasmids are strongly and positively correlated with IS26 copy numbers. Such correlations were further validated by *Escherichia coli* plasmids collected from human and pig wastewater. We propose an integron-IS26-ISCR (3I) framework with four MDR plasmid evolutionary stages of increasing ARG accumulation potential. The “3I” framework refines predictions of MDR plasmid emergence and spread, providing universal genetic markers for cost-effective surveillance.

## 1 Introduction

The emergence and spread of multidrug resistant (MDR) “superbugs” drive increased morbidity, mortality, and costs ^1–3^. Plasmids are the major carriers of antibiotic resistance genes (ARGs) in bacteria in human guts, drinking water, and food production chains ^4–6^. MDR plasmids are especially efficient in spreading ARGs in bacterial communities ^7,8^. In the recent Yemen cholera outbreak, the evolutionary success of a *Vibrio cholerae* lineage was linked to the carriage of an IncC MDR plasmid ^9^. Plasmids can gain or lose functional genes like ARGs by smaller mobile genetic elements (MGEs) like transposons and integrons to allow bacterial cells quickly adapt to dynamic selective pressures ^10–13^. Though it is widely accepted that selective pressure can maintain MDR plasmids in bacterial communities ^12,14^, predicting the emergence and spread of MDR plasmids remains a formidable challenge ^15^. The mechanisms governing the evolutionary transition of specific plasmids toward complex MDR architectures remain poorly understood.

Integron is one MGE whose presence is strongly linked to MDR bacteria in both anthropogenic and natural environments ^16–18^. Integrons can capture ARGs into their cassette arrays by *attC*-mediated site-specific recombination ^19^. However, the frequent co-existence of integrons with ARGs lacking *attC* sites, such as those conferring resistance to macrolides and tetracyclines, suggests additional mechanisms of plasmid-borne resistance ^18,20,21^. IS26 elements complement this limitation by facilitating sequence-independent mobilization and iterative tandem stacking ^22,23^. IS26 elements often truncate integrons, causing the prevalent formation and mobilization of partial integrons which may offer superior fitness over full-length architectures ^24,25^. Furthermore, ISCR elements utilize rolling-circle transposition to recruit non-cassette ARGs ^26^. In large-scale genomic studies over the past decade, ISCR elements have become increasingly underrepresented ^27–29^, likely due to their frequent incorporation into complex genetic contexts ^30^.

The characterization of integrons and insertion sequences in plasmids was challenging because short-read next-generation sequencing has limitations in identifying repeated regions. The development and application of long-read third-generation sequencing produces abundant high-quality plasmid assemblies, making it possible to investigate the plasmid ARG arrangement in high resolution. In this study, we characterized the combined effects of integrons, IS26, and ISCR elements on plasmid-level ARG accumulation. We propose a “3I” framework integrating observed stepwise ARG accumulation with established MGE mechanisms. Our findings can inform future predictive modeling and surveillance strategies for MDR emergence and dissemination.

## 2 Results

### 2.1 The increased ARG load on integron-carrying plasmids was not solely due to ARGs located in integron cassette arrays

Overall, 1,102 among the 3,211 circular plasmids retrieved from PLSDB RefSeq carried at least one ARG. Among these 1,102 ARG-carrying plasmids, 419 were integron-positive (i.e., at least one integron-associated structure was identified on the plasmid, including complete integron, In0, or CALIN element), and 683 were integron-negative (**Supplementary Table 1**). The number of ARGs carried by integron-positive plasmids were significantly higher than integron-negative plasmids, with medians of 9 and 2, respectively (p < 0.0001) (**Extended Data Fig. 1**). This trend was consistently observed across different incompatibility groups, bacterial host genera, and plasmid sizes (**Fig. 1a, b, c**). After excluding the integron-borne cassette ARGs, integron-positive plasmids still carried significantly more ARGs than integron-negative plasmids (p < 0.0001) (**Supplementary Note 1, Extended Data Fig. 2**). The classes of ARGs located inside and outside integrons varied on integron-positive plasmids. Aminoglycoside resistance genes were the most predominant both inside and outside integrons. ARGs conferring resistance to trimethoprim, quaternary ammonium, and rifamycin tended to distribute inside integrons, while those conferring resistance to beta-lactam, tetracycline, macrolide, and quinolone were more frequently located outside integrons (**Fig. 1d**). The significantly higher non-cassette ARGs on integron-positive plasmids and the varied distributions of ARG classes suggest the presence of integrons was not the only factor that contributed to plasmid ARG accumulation.

**Fig 1.**
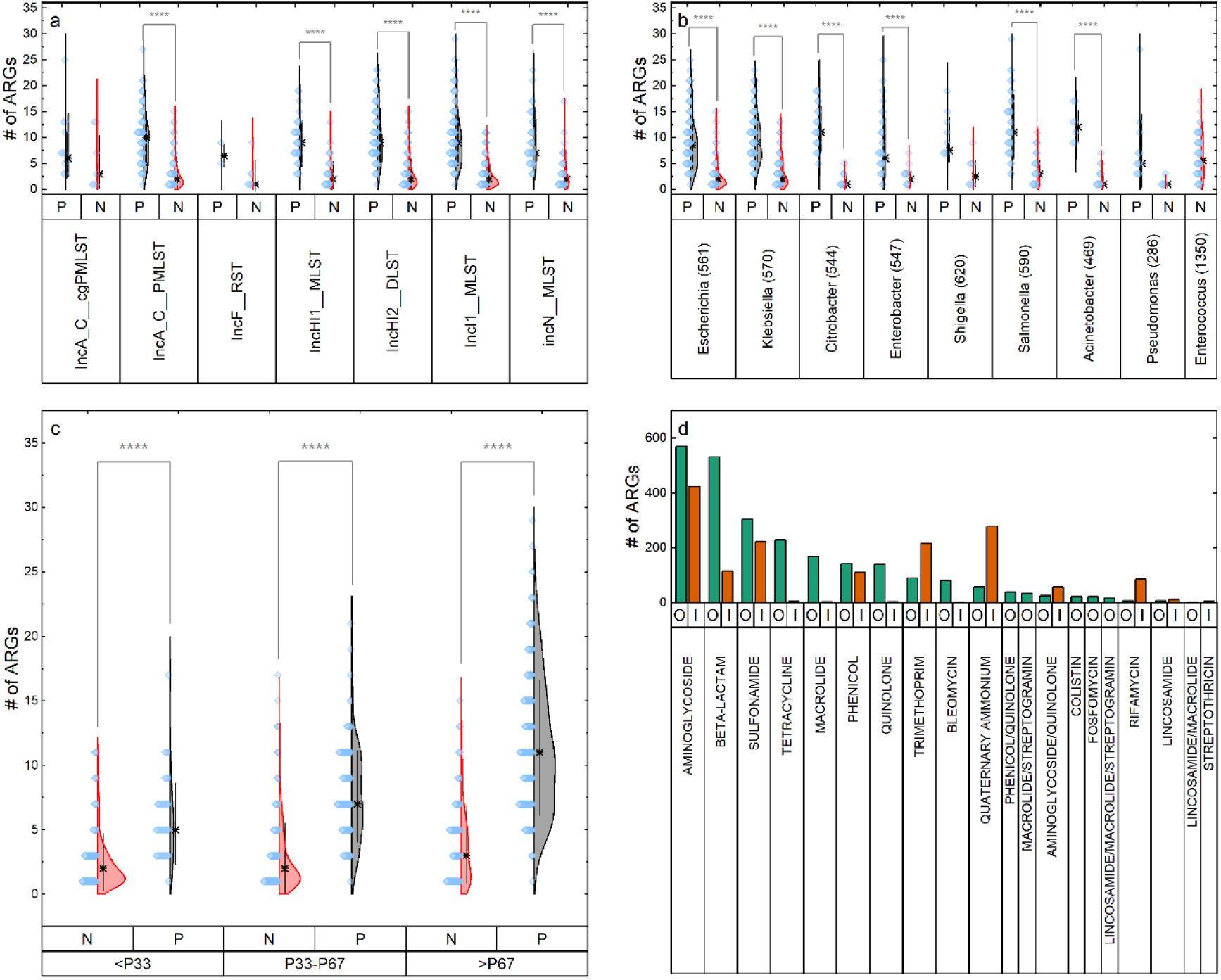
The presence of integrons led to significant increase in ARG counts in plasmids, with varied ARG classes inside and outside integron regions. a) Kruskal-Wallis ANOVA followed by Dunn’s Test for pairwise comparisons showed that the ARG counts on integron-positive plasmids were significantly higher than integron-negative plasmids across incompatibility groups, including IncA (n = 18 for IncA_C_cgPMLST, n = 258 for IncA_C_PMLST), IncH (n = 77 for IncHI1_MLST, n = 250 for IncHI2_DLST), IncI (n = 352), and IncN (n = 138) (p < 0.0001). There were too few IncF plasmids (n = 9) to generate significance. b) Kruskal-Wallis ANOVA followed by Dunn’s Test for pairwise comparisons showed that the ARG counts on integron-positive plasmids were significantly higher than integron-negative plasmids across bacterial host genera, including *Escherichia* (n = 337), *Klebsiella* (n = 280), *Citrobacter* (n = 34), *Enterobacter* (n = 57), *Salmonella* (n = 84), and *Acinetobacter* (n = 37) (p < 0.0001). *Shigella* (n = 18) and *Pseudomonas* (n = 17) had too few plasmids to generate significance. There was no integron-positive plasmid hosted by *Enterococcus* in this dataset. The numbers in the brackets next to genera names are the taxonomy IDs of the corresponding genera. c) Kolmogorov-Smirnov Test showed that integron-positive plasmids contain significantly more ARGs than integron-negative plasmids in all size ranges (p < 0.0001). The asterisks mean the median values of the number of ARGs in each group divided by the 33rd and 67th percentile cutoffs (P33 = 58,125 bp, P67 = 118,444 bp) (<P33: n = 42 for integron-positive, n = 321 for integron-negative; P33-P67: n = 130 for integron-positive, n = 245 for integron-negative; >P67: n = 247 for integron-positive, n = 117 for integron-negative). d) The classes of ARGs distributed inside (I, orange) or outside (O, green) of integrons on integron-positive plasmids.

### 2.2 With integron presence, the number of ARGs on plasmids was strongly correlated with the number of IS26 elements

Because IS26 elements contribute to the mobility of integrons and a wide variety of ARGs^24,25,31^, here, we determined the correlation between the ARG load and the IS26 copies in the 1,102 ARG-carrying plasmids with the presence or absence of integrons. As expected, integron-positive plasmids were more likely to harbor IS26 elements than in integron-negative plasmids (**Fig. 2a**). As shown in **Fig. 2b**, with the presence of both integrons and IS26 elements, the plasmids reached the significantly highest ARG load, with a median value of 10. The median ARG load dropped to 6 for integron-only plasmids and dropped to 2 for IS26-only plasmids. Without the presence of integrons or IS26 elements, the median ARG load reached the significantly lowest level of 1. These findings suggest a potential synergy between integrons and IS26 elements, as their combined presence resulted in a median ARG load (10) that exceeded the additive sum of their individual effects (6 and 2, respectively).

**Fig 2.**
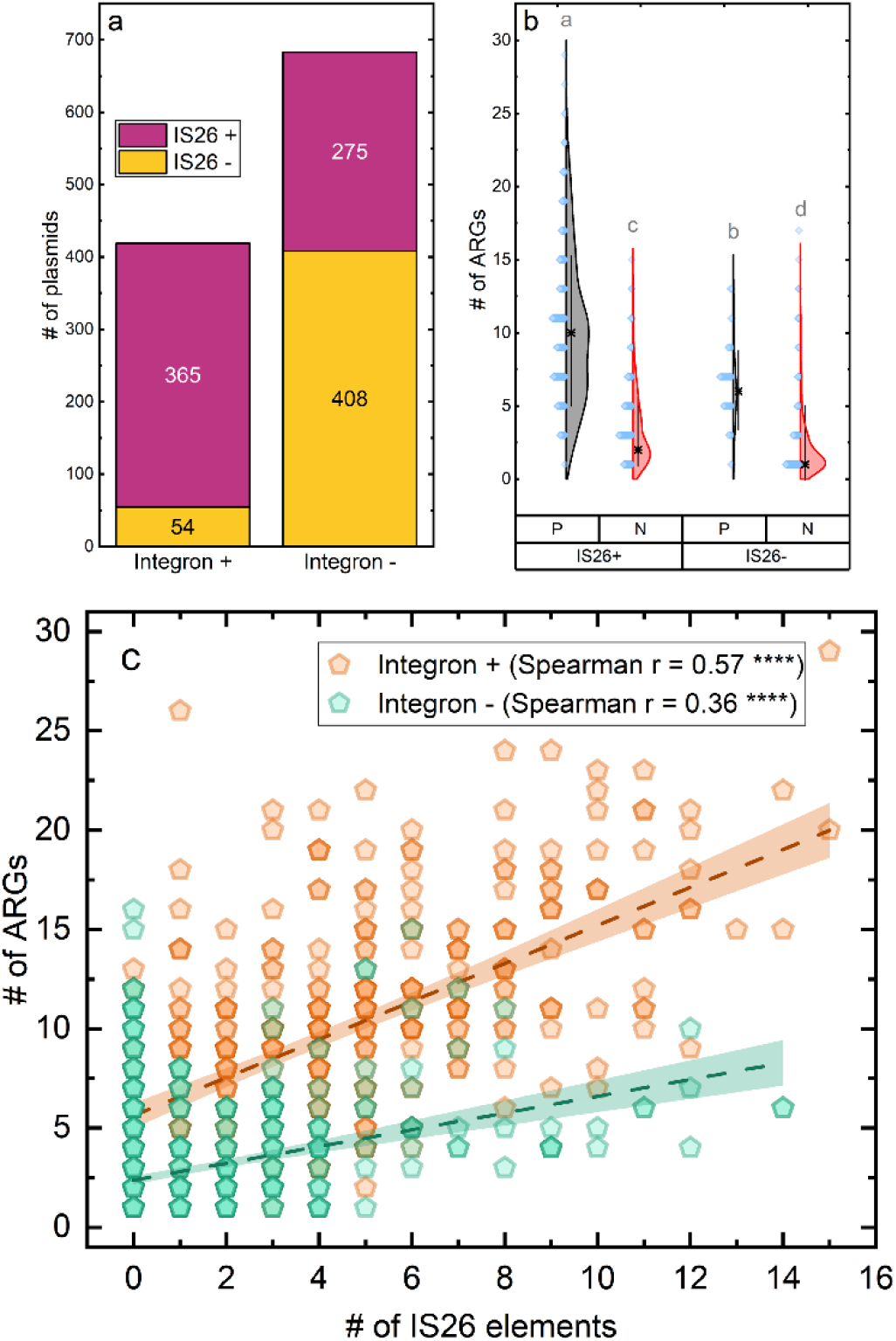
Characterization of plasmid ARG load impacted by integrons and IS26 elements. a) More integron-positive plasmids contain IS26 insertion sequences than integron-negative plasmids. b) Kruskal-Wallis ANOVA followed by Dunn’s Test for pairwise comparisons showed that the number of ARGs on the plasmids with or without IS26 or integrons present (P: integron-positive; N: integron-negative). The median numbers of ARGs for integron-positive IS26-positive (n = 365), integron-positive IS26-negative (n = 54), integron-negative IS26-positive (n = 275), and integron-negative IS26-negative (n = 408) plasmids were 10, 6, 2, and 1, respectively (p < 0.01). c) The number of ARGs versus the number of IS26 elements on integron-positive and integron-negative plasmids (****: p < 0.0001).

Ordinary Least Squares (OLS) regression analysis indicated that the synergy between integrons and IS26 elements explained nearly 60% of the variance in plasmid ARG load (p < 0.0001, R^2^ = 0.598) (**Supplementary Data 1**). To account for the non-normal distribution of residuals observed in the OLS model, a Robust Linear Model (RLM) was applied to ensure the stability of the estimates. A highly significant interaction was observed between the two MGEs (interaction coefficient = 0.58, p < 0.001). This interaction effectively doubled the efficiency of ARG accumulation from 0.42 ARGs per IS26 copy in integron-negative plasmids to 1.00 ARG per IS26 copy in integron-positive plasmids (**Fig. 2c**). The positive correlation between the number of ARGs and IS26 copy number was stronger for integron-positive plasmids than integron-negative plasmids, regardless of plasmid size (Spearman correlation coefficients of 0.57 versus 0.36; Spearman partial correlation coefficients of 0.48 versus 0.32; p < 0.0001 for all) (**Table 1**). Notably, after controlling for IS26, the correlation between plasmid size and ARG load in integron-negative plasmids was statistically significant but negligible (Spearman partial correlation coefficient of 0.08, p < 0.05), indicating that integrons and IS26 elements are the main drivers of ARG accumulation on plasmids, and simple size increases are insufficient to drive this process as effectively as the two MGEs.

**Table 1.** The Spearman correlation coefficients (out of brackets) and Spearman partial correlation coefficients (in brackets) among plasmid size, number of IS26 elements, and number of ARGs for integron-positive and integron-negative plasmids.

|  | **Plasmid size & #**<br>**of IS26 elements** | **Plasmid size & #**<br>**of ARGs** | **# of IS26 elements &**<br>**# of ARGs** |
| --- | --- | --- | --- |
| **Integron +** | 0.38****(0.13**) | 0.49****(0.37****) | 0.57****(0.48****) |
| **Integron -** | 0.35****(0.31****) | 0.20****(0.08*) | 0.36****(0.32****) |

(\*: $p < 0.05$ ; \*\*: $p < 0.01$ ; \*\*\*\*: $p < 0.0001$ )

### 2.3 The integron-IS26 synergy in ARG accumulation was also observed in the wastewater-borne plasmids collected from designated sampling sites

To ensure that the integron-IS26 synergy in plasmid ARG accumulation was not merely a byproduct of antibiotic-driven co-selection, we analyzed 32 *E. coli* plasmids isolated from designated wastewater sampling sites (**Supplementary Table 2**), including: human community-level raw sewage from two adjacent towns (Town A and Town B); anonymized, de-identified human clinical samples from a hospital in Town A; and wastewater from a pork processing center in Town B (**Fig. 3a**). Like the database-derived plasmids, wastewater-borne integron-positive plasmids carried significantly more ARGs than integron-negative plasmids, with medians of 9 and 1, respectively (p < 0.0001) (**Fig. 3b**). Co-existence of integron-positive and integron-negative ARG-carrying plasmids were observed in Town A, Town B, and the pork processing plant, with varied frequencies (**Fig. 3c**). Within the same sampling site, like Town B and pork processing plant, integron-positive plasmids carried significantly more ARGs than integron-negative plasmids (p < 0.05) (**Fig. 3d**), and most ARGs on integron-positive plasmids were located outside integron regions (**Extended Data Fig. 3**). Consistent with database-derived plasmids (**Fig. 2c**), the 11 wastewater-borne integron-positive plasmids also exhibited higher IS26-correlated ARG accumulation efficiency than the 21 integron-negative plasmids (**Fig. 3e**). To conclude, while selective pressure favors the enrichment of MDR plasmids in sites like pork processing plant (**Fig. 3c**), our results suggest that MGEs like integrons and IS26 elements act as the primary mechanistic drivers for ARG accumulation within these plasmids (**Fig. 3d, e**).

**Fig 3.**
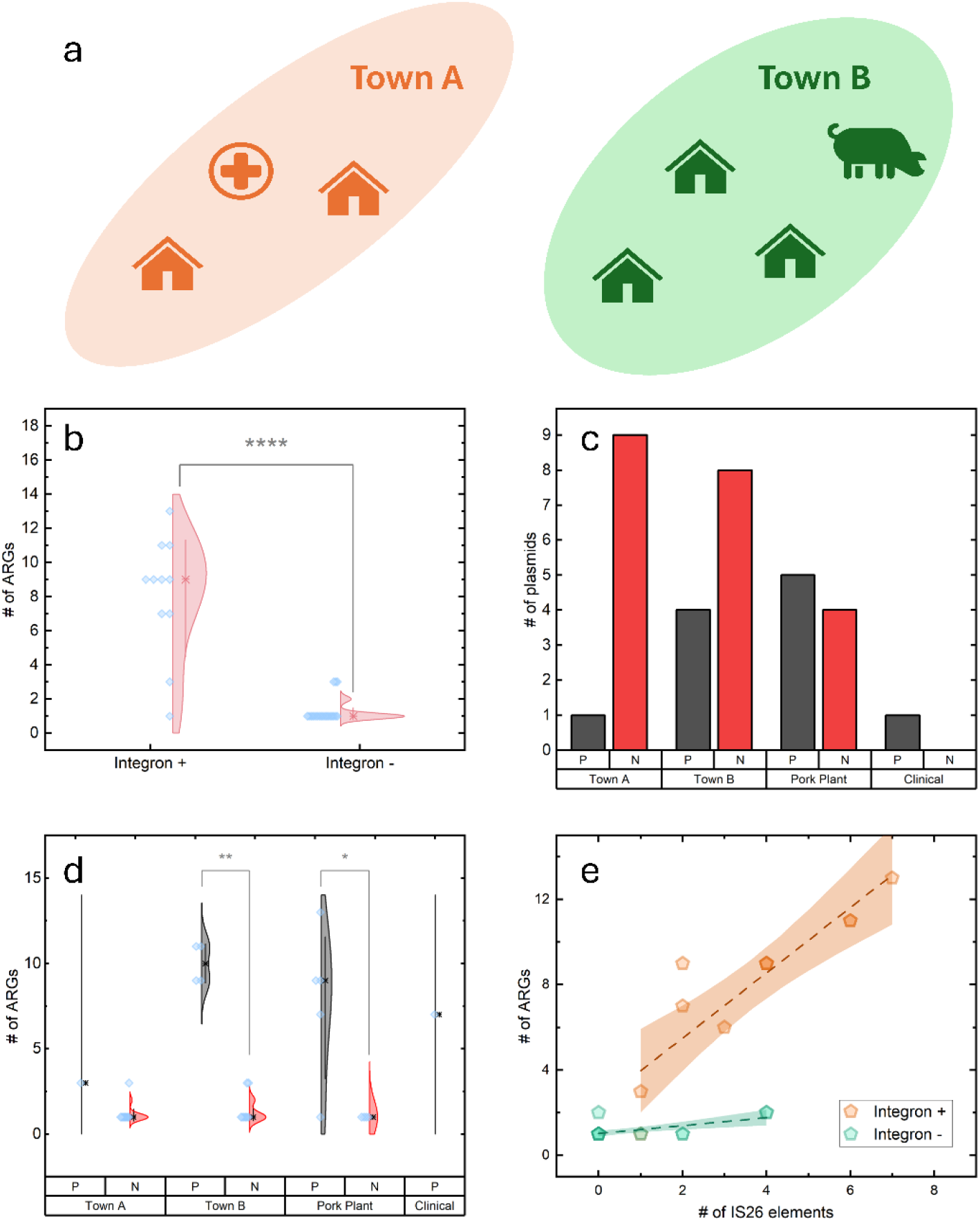
The number of ARGs, integron-positive plasmids, and integron-negative plasmids in environmental *E. coli* samples collected in this study. a) The sampling sites in Town A and Town B. Town A sampling sites included two human communities and a hospital, while Town B sampling sites included three human communities and a pork processing center. b) Kolmogorov-Smirnov Test showed that the number of ARGs on integron-positive plasmids (n = 11) was significantly higher than the number of ARGs on integron-negative plasmids (n = 21) in the *E. coli* strains collected in this study (p < 0.0001). c) The number of integron-positive and integron-negative ARG-carrying plasmids collected from each sampling site (Town A: 10; Town B: 12; Pork Plant: 9; Clinical: 1). d) For Town B and pork plant, the number of ARGs on integron-positive plasmids was significantly greater than those on integron-negative plasmids (p < 0.01 for Town B; p < 0.05 for pork plant). Town A and clinical samples had too few integron-positive plasmids to do statistical test. e) The number of ARGs versus the number of IS26 elements on the 32 integron-positive (orange) or integron-negative (green) ARG-carrying *E. coli* plasmids.

### 2.4 ISCR elements, whose presence is strongly associated with the presence of integrons or IS26 elements, are linked to the highest ARG accumulation level on plasmids when co-existing with IS26 elements and integrons

We further investigated if other MGEs were involved in the integron-IS26 driven plasmid ARG accumulation. ISCR elements, specifically ISCR1 and ISCR2, were examined because they are frequently embedded within integron-IS26 frameworks ^26^. The plasmids with all three MGEs (i.e., integron, IS26, and ISCR) present possessed the highest ARG count among all groups (**Fig. 4a**). Most ARGs on these plasmids were non-redundant (**Extended Data Fig. 4**). Notably, the median ARG count followed a downward trend as MGE diversity diminished, highlighting the synergistic role of integrons, IS26, and ISCR elements in building complex MDR profiles. When there were no integron, IS26, or ISCR elements present, the median number of ARGs on a plasmid reached the lowest level.

**Fig 4.**
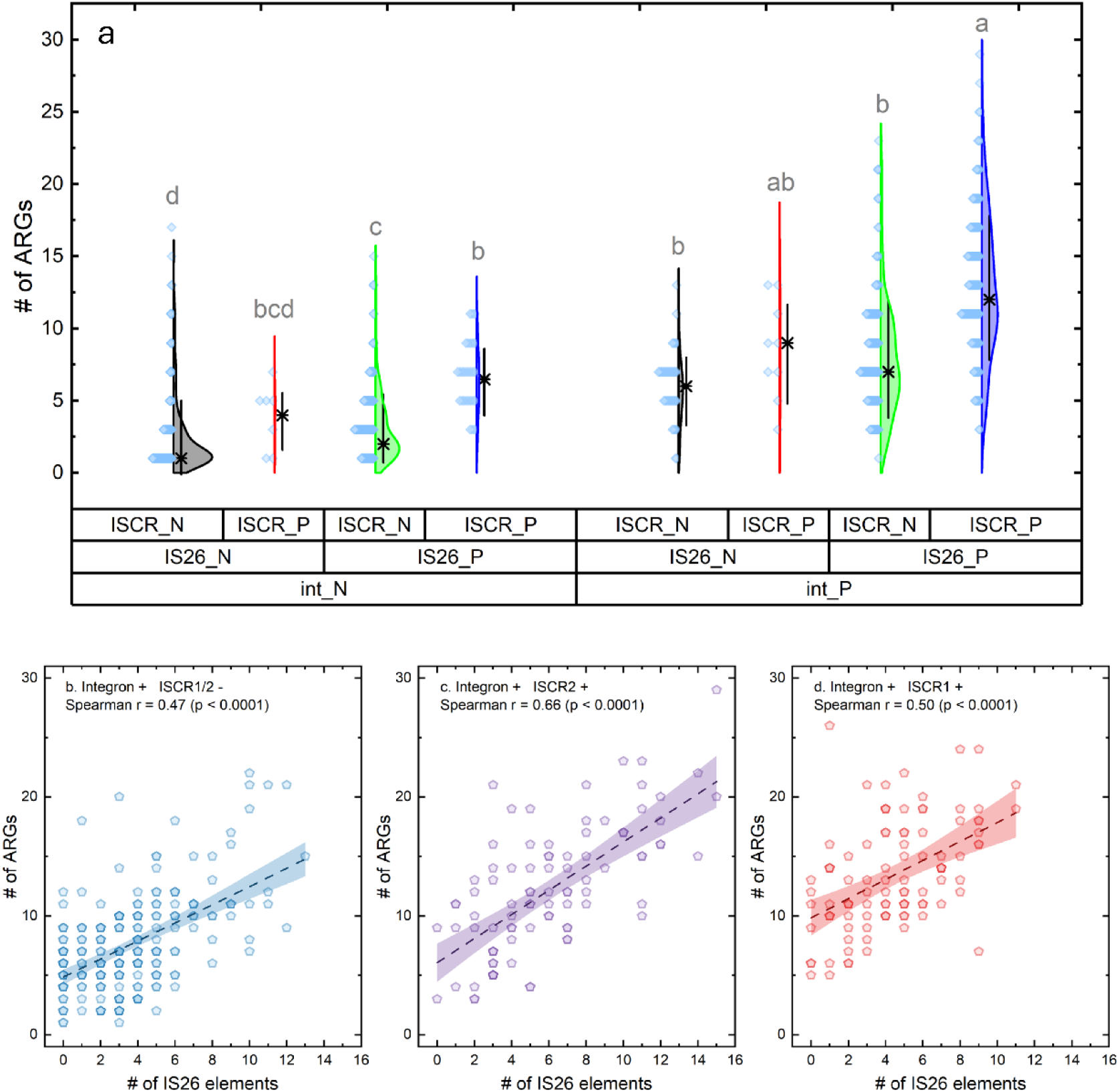
Characterization of plasmid ARG load impacted by different combinations of integrons, IS26, and ISCR elements. a) The number of ARGs on plasmids with the presence or absence of integrons, IS26 elements, and ISCR elements. Significant differences (p < 0.05) were tested by Kruskal-Wallis ANOVA with Dunn’s Test for pairwise comparisons and are indicated by distinct letters in the figure. The plasmids with all three MGEs present (n = 166) possessed the highest ARG count among all groups. For the plasmids with integron and ISCR presence but IS26 absence, the data points were too few to generate significance (n = 9). b) The correlation between the number of ARGs and the number of IS26 elements on integron-positive plasmids without ISCR1 or ISCR2 elements. When there were no ISCR1 or ISCR2 elements on the plasmid, the Spearman correlation coefficient between the number of ARGs and the number of IS26 elements was 0.47 (n = 244, p < 0.0001) c) The correlation between the number of ARGs and the number of IS26 elements on integron-positive plasmids with ISCR2 elements. When integron co-existed with ISCR2 elements, the Spearman correlation coefficient increased to 0.66 (n = 91, p < 0.0001). d) The correlation between the number of ARGs and the number of IS26 elements on integron-positive plasmids with ISCR1 elements. When integron co-existed with ISCR1 elements, the Spearman correlation coefficient was 0.50 (n = 99, p < 0.0001).

To elucidate how the distinct genetic context preferences of ISCR1 and ISCR2 elements influence plasmid ARG recruitment, we compared IS26-mediated ARG accumulation patterns for integron-positive plasmids carrying ISCR1, ISCR2, or neither (**Fig. 4b, c, d**). With the presence of both integrons and ISCR2 elements, the positive correlation between ARG counts and IS26 copies was the strongest (Spearman r = 0.66, p < 0.0001). The co-existence of integron and ISCR1 elevated the baseline ARG count, regardless of the IS26 copy number (RLM intercept shift of 4.39, p < 0.001), while the co-existence of integron and ISCR2 led to a higher recruitment rate of ARGs per IS26 copy (RLM slope difference of 0.34, p = 0.007) (**Supplementary Data 1**). Interestingly, the presence of ISCR elements was scarce without IS26 presence (n = 7 for integron-negative and n = 9 for integron-positive) (**Fig. 4a**), and the presence of integrons also was highly correlated with IS26 presence (**Extended Data Table 1**). These correlations suggest there might be an underlying evolutionary logic that maintains the association among the three MGEs.

## 3 Discussion

Here, we propose a structural, integrative model for MDR plasmid evolution, in which integrons serve as structural scaffolds guiding plasmid evolution toward multidrug resistance; ISCR elements contribute to the expansion of non-cassette ARG content; and varied copy numbers of IS26 elements provide structural flexibility that facilitates ARG accumulation and rearrangement. Based on the plasmid ARG accumulation patterns observed in this study, here, we propose a “3I” framework to describe the evolution of MDR plasmids involving integrons, IS26, and ISCR elements, as shown in **Fig. 5**. This framework includes four stages of plasmid evolution with increased types of MGEs. At stage 0, the plasmid is free of integrons, IS26 elements, or ISCR elements, with very few ARGs present. Then, at stage I, the plasmid has either the first integron inserted by MGEs like Tn21 or Tn402 ^19,23^, or the first IS26 inserted by random insertion. Independent ISCR element presence is excluded from **Fig. 5**, as they rarely occur without the other two MGEs (7 in 1,102) (**Fig. 4a**). At stage II, IS26 elements and integrons start to co-exist in plasmids, either by IS26 insertion into an integron-carrying plasmid, or by IS26 mobilizing an integron into a plasmid. At this stage, though the median ARG count was not significantly greater than the plasmids with only integron presence, the maximum ARG capacity of a plasmid increases (**Fig. 4a**). IS26 elements also can delete IS26-flanked gene cargos from the DNA backbone ^22^. At stage II, by specifically activating this deletion reaction targeting the IS26-flanked integron region, it would be possible to invert the plasmid back to stage I. At stage III, integrons, IS26 elements, and ISCR elements co-exist in the plasmid, making both the median number of ARGs and the maximum ARG capacity increase (**Fig. 4a**).

**Fig 5.**
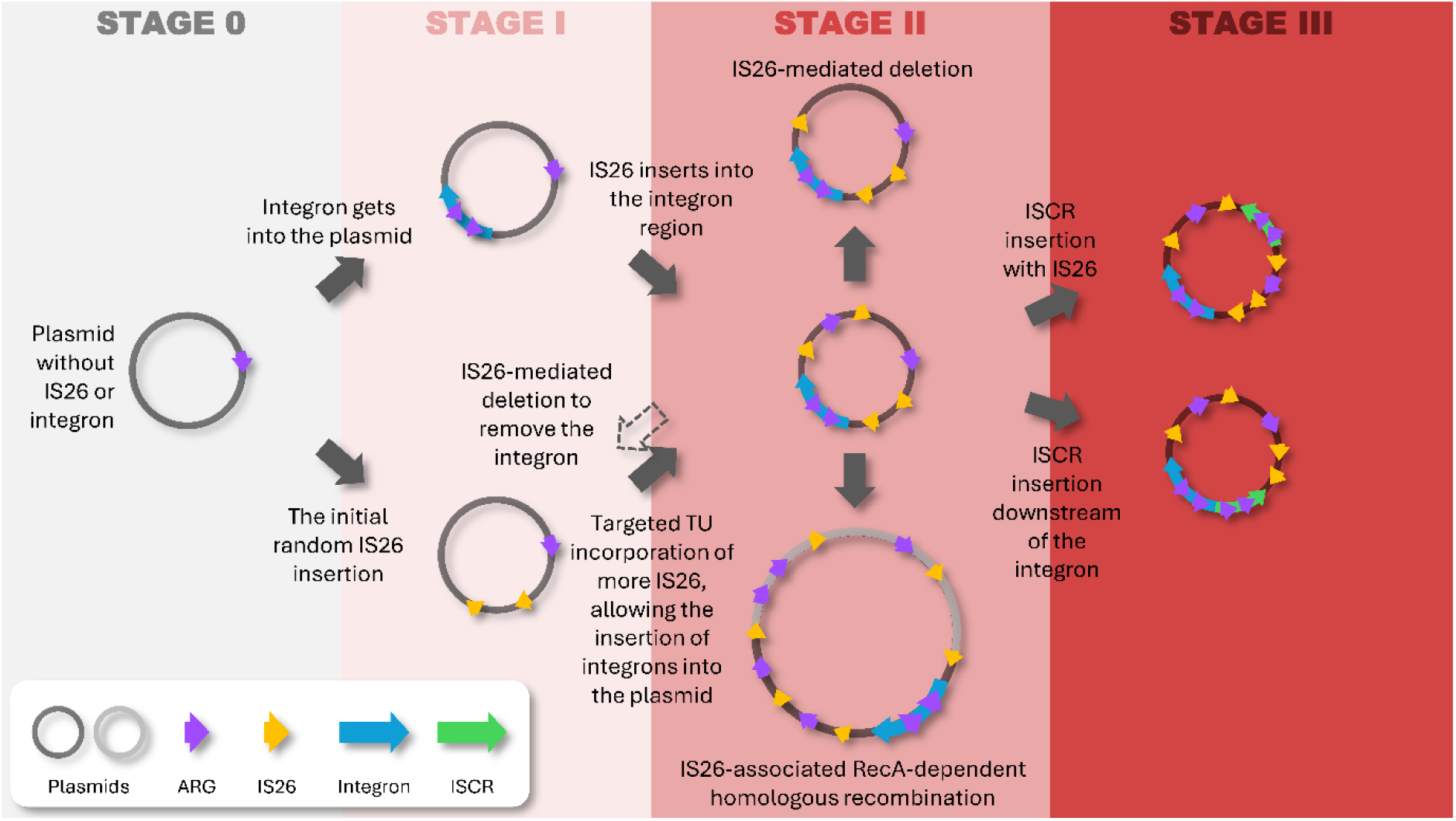
Proposed “3I” framework for integron-IS26-ISCR-associated stepwise ARG accumulation on plasmids. The dashed arrow suggests a potential intervention strategy to turn stage II plasmids back to stage I.

It is important to highlight that the “3I” framework here is more about the plasmid ARG accumulation potential, instead of the actual number of ARGs carried by a plasmid. In this study, we observed strong positive correlations between the copy numbers of ARGs and IS26 elements on integron-positive and ISCR-positive plasmids (Spearman correlation coefficients varied from 0.47 to 0.66, p < 0.0001) (**Fig. 2c, 4b, 4c, 4d**). These strong correlations suggest that though the MGE combination sets the capacity for ARG acquisition, the realized extent of accumulation is driven by IS26 copy number. IS26 elements have a distinctive property that they tend to insert adjacent to existing IS26 elements by translocatable unit (TU) incorporation, with efficiency around 60-fold higher than random insertion ^23^. Such a property can allow the fast iterative stacking of IS26 elements with bounded DNA segments once the first IS26 element is inserted into the plasmid ^32^. In addition, the conserved sequences on IS26 elements can allow RecA-dependent homologous recombination, resulting in the formation of a large recombinant plasmid with diverse ARGs originated from the two involved plasmids (**Supplementary Note 2, Extended Data Fig. 5**), though such homologous recombination is far less efficient than IS26 TU incorporation ^33^. Though ARGs are the major cargo genes mobilized by IS26, heavy metal resistance genes like *silE* and *copM* were found flanked by IS26 elements in NZ_CP021103.1. The decreased Spearman correlation coefficient of 0.36 for integron-negative plasmids (**Fig. 2c**) suggests that IS26 elements also contribute to the mobilization of other functional genes, especially without integron presence.

In addition to the fast iterative stacking property of IS26 elements, DNA segments that are bound by two IS26 elements can also be deleted from the plasmid under certain conditions ^22^. The deletion mechanism of IS26 elements can allow bacteria to quickly adapt to the fluctuating environmental pressures. The potential deletion events were also observed in seven of the 10 plasmids as shown in **Supplementary Note 2** and **Extended Data Fig. 5**. In this study, the strong positive correlation between the ARG and IS26 copy numbers on integron-positive and ISCR-positive plasmids may not only be due to the iterative stacking of IS26 elements, but also the deletion events that remove the flanked ARGs from the plasmid backbone as well as reduce the IS26 copy number.

As antibiotic drug discovery gets increasingly challenging, the focus of antimicrobial resistance (AMR) research strategically shifted toward the evolutionary mechanisms of bacterial adaptation. Based on the findings of this study, inducing the deletion activity of IS26 elements during or after antibiotic usage may be a promising way to mitigate the rapid development of multidrug resistance in the future. In **Fig. 4a**, though the median numbers of ARGs were significantly different among plasmids with different MGE combinations, each group had a certain number of plasmids with five or fewer ARGs, mostly with few IS26 copies as well, suggesting the possibility of reversing the multidrug resistance on plasmids at all plasmid evolution stages by reducing the IS26 copy numbers. Unlike gene-editing strategies like CRISPR that require the delivery of large molecules into bacteria cells, inducing the deletion activity of IS26 elements may only require the delivery of small molecules that can interact with the Tnp26 transposase which already exist in many MDR bacteria ^34^.

While IS26 mediates plasmid ARG accumulation through iterative stacking, integrons and ISCR elements act as foundational platforms for the compact recruitment of ARGs. The gene acquisition mechanism of integrons requires site-specific recognition of *attC* structures ^19^. Though *attC* sites can be found in ARGs conferring resistance to aminoglycoside and trimethoprim ^35,36^, genes conferring resistance to quinolone, tetracycline, and macrolide usually do not include *attC* sites, and are usually located outside integrons (**Fig. 1d**). The recruitment of these non-cassette ARGs can be mediated by ISCR elements through a rolling-circle-mediated mechanism ^26^. With the ARGs acquired by integrons and ISCR elements, IS26 elements can mobilize and rearrange these genes in a site-independent manner ^22^.

The mobilization of integrons often is associated with IS26 elements ^22^. IS26 elements frequently are inserted into integrons, causing the loss of the 3’ CS region when integrons move ^24,25^. Such movement of partial integrons also was described in **Supplementary Note 2**. With the *intI* and *attI* regions intact, the loss of the 3’ CS region would not inhibit IntI-mediated *attI* site-specific recombination ^19^. The insertion of an IS26 element into an integron can also serve as a hotspot for future insertion of IS26-flanked mobile gene cargos ^37^. Though previous study found nearly equal number of complete integron and CALIN in bacterial genomes ^38^, most integron-associated structures on the plasmids collected in this study were complete integrons or In0 with intact *intI* and *attI* regions, instead of CALIN which lack the gene acquisition functions (**Extended Data Fig. 6**). The high frequency of complete integrons and In0 instead of CALIN on plasmids are in alignment with the active plasmid gene exchange driven by integrons identified in previous studies ^28,29^. Though relatively rare, the structure of CALIN flanked by two either directly or inversely oriented IS26 elements were also observed in NZ_CP041112.1, NZ_CP058835.1, NZ_CP095650.1, and NZ_KX246266.1 (**Extended Data Fig. 7**), suggesting that both integrase modules and gene cassette arrays of integrons can be mobilized or rearranged by IS26 elements.

Despite the shared rolling-circle-mediated mechanisms, ISCR1 was associated with a higher baseline ARG load in the absence of IS26, while ISCR2 appeared to potentiate a more rapid expansion of the ARG load as IS26 copies increased (**Fig. 4c, d**). The different mobilization preferences between the two ISCR elements may be the underlying driver of the difference in ARG-IS26 correlation patterns. ISCR1 often serves as a functional extension of class 1 integron by mobilizing resistance genes that lack *attC* sites, while ISCR2 can operate independently of the integron framework and can be mobilized by MGEs like IS26 ^26^. Unlike integrons and IS26 elements, ISCR elements were rarely explicitly addressed in recent large-scale ARG-MGE studies ^28,29^, likely because their presence was highly correlated with integrons and IS26 elements (**Extended Data Table 1**). Because the co-existence of integrons, IS26, and ISCR elements contributed to the highest level of ARG accumulation on plasmids (**Fig. 4a**), it is worthwhile revisiting some large-scale bioinformatics studies on ARG-MGE correlations to understand the underrepresented contributions of ISCR elements.

Furthermore, the evolutionary success of the combination of integrons, IS26, and ISCR elements was demonstrated during the cholera outbreak that occurred in Yemen from 2016 to 2019 ^9^. A pseudo-compound transposon containing all these three MGEs with eight ARGs conferring resistance to various classes of antibiotics, which was likely created through multiple independent insertion events, was disseminated with an IncC plasmid, and the ratio that the dominant lineage of *Vibrio cholerae* (*Vc*H.9) carried this specific MDR plasmid increased from 6.7% (6/89) in 2018 to 100% (151/151) in 2019 ^9^. Researchers also called for real-time surveillance and diagnostic strategies to differentiate low-, middle-, and high-risks during the emergence and spread of MDR strains ^9^.

Currently, surveillance for MDR strains occurs at a relatively late stage after a MDR strain is uncovered, sometimes focusing on the incompatibility groups of the identified MDR plasmid ^39^. Our results suggest that incompatibility groups do not have as significant contributions to the plasmid ARG load as MGEs like integrons, IS26, and ISCR elements (**Fig. 1a, 4a**). Though integrons have long been considered as indicators of multidrug resistance ^19,40,41^, the “3I” framework proposed in this study establishes integrons, IS26, and ISCR elements as pivotal genetic markers that allow more precise prediction of MDR risks before the fast accumulation of ARGs. By understanding the correlations between the plasmid ARG load and the presence of the three MGEs, detecting integrons, IS26, and ISCR elements by qPCR or biosensors would allow the prompt identification of what stage the MDR plasmids have evolved. Because the plasmid ARG load is strongly correlated with the IS26 copy number (**Fig. 2c, 4b, 4c, 4d**), a further step of quantifying IS26 elements could allow the estimation of the ARG load. With such large-scale cost-effective qPCR screenings, whole-genome sequencing can be applied specifically to the strains with high MDR risks for a more accurate and targeted identification of the ARGs that are carried by the MDR plasmids. Such a surveillance framework could be particularly effective in resource-limited settings, such as during the Yemen cholera outbreak.

To conclude, in this study, we characterized the hierarchical “3I” framework for stepwise ARG accumulation on plasmids associated with the presence of integrons, IS26, and ISCR elements. The co-existence of the three MGEs was linked to the highest ARG accumulation level on plasmids. Despite the insights provided by this study, several limitations should be noted. First, other MGEs, including IS903 and ISEcp1, may also contribute to MDR plasmid evolution at different functional scales. Second, the public dataset used here may overrepresent plasmids from clinical environments characterized by elevated antibiotic selective pressures. Third, because this study is based on genomic sequences, ARG expression and phenotypic resistance could not be directly assessed. Notably, though no integron-positive *Enterococcus* plasmids were identified in this study, and it had long been considered that gram-positive bacteria like *Enterococcus* would not harbor integrons, recent studies started to report the presence of integrons in some *Enterococcus* strains ^42,43^. The newly emerged integron-positive *Enterococcus* strains suggested that a similar combination of MGEs for ARG accumulation may soon occur in gram-positive bacteria, though the role of IS26 may be replaced by other insertion sequences like IS1216 with similar functions but more frequently identified in gram-positive bacteria genomes ^44^. The positive correlation between IS26 element copies and ARG copies on integron- and ISCR-positive plasmids identified in this study can guide the future development of IS26-deletion-driven AMR mitigation strategies. We also identified the underrepresented role of ISCR elements contributing to the plasmid ARG accumulation, suggesting a need for a better characterization of this MGE family. Ultimately, the “3I” framework proposed here provides a robust blueprint for deciphering hierarchical ARG acquisition, paving the way for advanced MDR predictive models and precision surveillance strategies.

## 4 Methods

### 4.1 Plasmid sequence retrieval and annotation

The sequences of the plasmids analyzed in this study were retrieved from PLSDB (data version: 2024_05_31_v2) ^45^. 3,211 circular RefSeq plasmid sequences with annotated incompatibility groups were downloaded based on their accessions (**Supplementary Table 3**). The genes on these plasmids were annotated by BV-BRC Genome Annotation using annotation recipe “Bacteria / Archaea” and taxonomy name “Bacteria (2)” ^46^. After annotation, 1,102 plasmids had at least one ARG annotated (**Supplementary Table 4**). Redundancy of the 1,102 ARG-carrying plasmids was assessed by FastANI with fragment length set to 1,000 ^47^. All 1,102 plasmids were retained to capture IS26-mediated micro-evolutions, given that plasmids with >99% average nucleotide identity (ANI) and coverage still possessed varying ARG profiles. The integron regions, including complete integron, clusters of *attC* sites lacking integron-integrases (CALIN), and integron-integrases lacking cassettes (In0), were identified by IntegronFinder 2.0 for these 1102 ARG-carrying plasmids ^48^. The ARGs located inside and outside of integron regions were determined by customized Python codes (**Supplementary Software 1**). ARGs with at least one base-pair overlapping with the integron region were identified as inside, while no overlapping was identified as outside. The number of unique ARGs versus the number of total ARGs on each plasmid was summarized by customized Python code (**Supplementary Software 1**). The presence of IS26 elements on the plasmids was determined by BLAST using the NCBI GenBank accession X00011.1 as the query sequence ^49^. To take into account the potential of RecA-dependent homologous recombination between homologous IS26 elements, plasmids with hits ≥ 80% identities and at least 500 bp matched lengths were considered to contain the IS26 elements. The presence of ISCR elements were determined by BLAST using NCBI GenBank accessions AAA92748.2 for ISCR1 transposase and UKC64023.1 for ISCR2 transposase as the query sequences ^50^. Because of the high amino acid similarity between ISCR1 and ISCR2 transposases, plasmids with hits ≥ 95% identities and at least 300 amino acid-matched lengths were considered to contain either ISCR1 or ISCR2 elements.

### 4.2 Statistical analysis on number of ARGs and MGE-carrying plasmids

To compare the numbers of ARGs on plasmids in different categories, normality test was first performed to determine whether the number of ARGs on plasmids were normally distributed. Two-sample Komogorov-Smirnov test was used to assess whether the distributions of the ARG numbers on integron-positive plasmids were significantly greater than those on integron-negative plasmids in pairwise. When more than two categories were compared with each other, Kruskal-Wallis ANOVA was used to assess whether the number of ARGs were distributed significantly differently among the categories. Chi-square tests with the calculation of odds ratios were conducted among the plasmids with presence of absence of integrons, IS26 elements, and ISCR elements, in order to understand the associations of the presence of the three MGEs. The statistical analyses were conducted on OriginPro 2024.

### 4.3 Case study of a representative cluster for plasmid evolution and comparative genomic alignment

To assess the dynamics of integrons and IS26 elements that contribute to different ARG patterns on plasmids, we obtained a plasmid lineage with highly similar backbones but diverse ARGs. First, the similarity of the 3,211 circular PLSDB RefSeq plasmids was assessed by calculating pairwise ANI using FastANI with minimum fraction of 0.5 and fragment length of 1000 ^47^. Cytoscape was used to plot networks based on the FastANI outputs ^51^. A cutoff of ANI > 99.5% was used to find plasmids that share highly similar sequences. Sixty-five plasmids that were clustered in the network with various numbers of ARGs and both integron-positive and integron-negative members were selected for a more detailed analysis on the ARGs on integron-positive and integron-negative plasmids. The core genome and pan genome of this plasmid cluster were extracted by PIRATE ^52^. Because no core genome was identified across the 65 plasmids, we plotted a phylogenetic tree using the pan genome data by FastTree and identified a clade with 11 plasmids that share highly similar pan genomes ^53^. Ten of the 11 plasmids adopted third-generation long-read techniques for sequencing. The core and pan genomes of these 10 plasmids were extracted by PIRATE ^52^. The core genome phylogenetic tree of the 10 plasmids was plotted by FastTree ^53^. The sequences of the 10 plasmids were aligned by Clinker using their annotated GenBank files ^54^.

### 4.4 Spearman correlation for plasmid sizes, ARG counts, and IS26 element copies

By dividing the 1,102 plasmids into integron-positive and integron-negative groups, the Spearman correlation coefficients among plasmid sizes, ARG counts, and IS26 element copies were calculated for each group using OriginPro 2024. Linear regression lines with 95% confidence intervals were added to scatter plots to aid the visualization of the monotonic trend. OLS and RLM regression analyses in Python package statsmodels were performed to compare the slopes and intercepts between linear regression lines (**Supplementary Software 1**). Specifically, RLM with Huber’s M-estimation was utilized to ensure regression stability against outliers. Spearman partial correlation coefficients were calculated using the equation below ^55^:

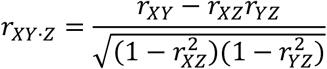

In the equation, r_XY·Z_ means the Spearman partial correlation coefficient between X and Y with the controlling variable as Z. r_XY_, r_XZ_, r_YZ_ mean the Spearman correlation coefficients between X and Y, between X and Z, and between Y and Z.

### 4.5 Environmental sample collection and processing

Weekly-composite wastewater samples from the five human communities in two towns (Town A and Town B) were taken from manholes using automatic water samplers (Teledyne ISCO). There is a pork processing center in Town B. Wastewater samples were collected at the pork processing center’s wastewater outlet. Each sample was stored in a sterile sampling bag during sampling and transportation (Fisherbrand). Once the sampling was completed, 20 mL of 2.5 M MgCl_2_ were added to each sample on-site for flocculation. Then, the sampling bags were sealed and transported to the laboratory in coolers filled with ice-packs within three hours. The samples were stored at 4 ℃ before the same-day pre-processing.

Samples were pre-processed in biosafety cabinets by filtering 10-50 mL well-mixed wastewater through 0.45 μm mixed cellulose ester (MCE) membrane filters (Millipore). A blank control was made by filtering 50 mL of molecular biology grade water. After filtering, each filter was cut into quarters. To maintain bacterial viability, each quarter was submerged in LB broth with 25% glycerol in a cryo-vial, pre-cooled at -20 ℃ for one hour, and stored at - 80 ℃ before *Escherichia coli* isolation (see below).

### 4.6 *E. coli* isolation and characterization

*E. coli* was isolated by Levine Eosin Methylene Blue (EMB) agar plates. Frozen filters were defrosted at 37 ℃ for 30 minutes. Then, the filters were placed onto EMB plates with the sediment side facing the agar. The plates were incubated at 37 ℃ for maximum of 24 hours. After incubation, colonies were re-streaked onto new EMB plates and incubated at 37 ℃ until round-shaped single colonies with dark purple color and green metallic sheen for identifying *E. coli* were present. Single colonies were re-streaked onto EMB agar plates an additional three times to eliminate the chance of getting mixed colonies. A blank control was included in every plate incubation step by streaking an EMB agar plate with phosphate buffered saline (PBS). A single presumed *E. coli* colony from each plate was enriched in cryo-vials with LB broth overnight at 37 ℃ with a shaking speed of 250 RPM to make frozen stocks. The blank control of the enrichment step was a cryo-vial with LB broth dipped by a clean inoculating loop. After enrichment, the bacteria culture was mixed with an equal volume of 50% glycerol and stored at -80 ℃ before downstream analyses.

16S rRNA gene Sanger sequencing was performed to verify that the solated strains were *E. coli*. Bacterial genome DNA was extracted by DNeasy Blood and Tissue Kit (QIAGEN). The bacteria cells for DNA extraction were obtained by re-streaking the frozen stock onto EMB plates once and enriching in LB broth as described above. All extraction steps followed the instruction provided by the manufacturer. All final DNA extracts were eluted with 200 μL AE buffer in the extraction kit. One extraction blank was inoculated with nothing and extracted using the abovementioned method. The DNA extracts were stored at −20 °C before the PCR of the 16S rRNA gene.

The reaction volume of PCR was 25 μL, containing 12.5 μL Taq 1x Master Mix (New England Biolabs), 1.25 μL forward primer(5’-TCGTCGGCAGCGTCAGATGTGTATAAGAGACAGCCTACGGGNGGCWGCAG-3’), 1.25 μL reverse primer(5’-GTCTCGTGGGCTCGGAGATGTGTATAAAGACAGCCTACGGGNGGCWGCAG-3’), 2 μL DNA sample and 8 μL nuclease-free water. The thermal cycle included a pre-denaturation stage of 95 ℃ for 15 min, a PCR stage with 40 cycles of 95 °C for 20s, 55 °C for 45s, and 68 °C for 45s, and a final extension stage at 68 °C for 5 min. After PCR, Exo-CIP™ Rapid PCR Cleanup Kit (New England Biolabs) was used to clean up the primers and polymerases. The PCR product was quantified by Qubit dsDNA BR Assay Kit (Thermo Fisher). Then, the DNA samples were submitted to a university core facility for Sanger sequencing. BLAST was conducted to compare the Sanger sequencing results to the nucleotide database on NCBI, and the bacteria strain was determined as *E. coli* if the top match of the BLAST result was *E. coli*.

### 4.7 *E. coli* long-read whole-genome sequencing and annotation

In addition to the environmental *E. coli* isolated in this study, we also received three *E. coli* strains isolated from de-identified human clinical samples from the healthcare facility located in Town A. The high-molecular weight (HMW) *E. coli* genome DNA was extracted by GenFind v3 (Beckman Coulter). The quantity and quality of the DNA extracts were determined by NanoDrop. The HMW DNA extracts were sheared by Megarupter 3 (Diagenode) at speed 40 before PacBio HiFi sequencing library preparation. Inhibitors were removed from the sheared DNA extracts by OneStep PCR Inhibitor Removal Kit (Zymo Research) followed by 1:1 bead cleaning using SMRTbell® cleanup beads (PacBio). PacBio HiFi sequencing libraries were made using the SMRTbell prep kit 3.0 (PacBio). The DNA yields of the libraries were determined by Qubit™ 1X dsDNA High Sensitivity (HS) Assay Kit (Invitrogen) and DNA fragment lengths were determined by AATI Fragment Analyzer by a university core facility before sequencing. Sequencing was conducted on Revio SMRTcell (PacBio) with 30-hour movie time by a university core facility. The sequencing reads were assembled *de novo* by Trycycler using Canu, Raven, and Flye as the assemblers ^56–59^. The whole-genome assemblies have been deposited in the NCBI BioProject database under accession number PRJNAXXXXXXX (Accession number will be provided upon publication). The *E. coli* genomes were annotated by BV-BRC Genome Annotation service. The integron regions were determined by IntegronFinder 2.0 ^48^.

