## Supplementary for "Synergistic interactions of mobile genetic elements shape the stepwise evolution of multidrug-resistant plasmids": Code_1_Summarize_integron_regions.py

```
import pandas as pd
import os

folder=FOLDER_PATH
file_list=[]
for root, dirs, files in os.walk(folder):
    for file in files:
        if file.endswith(".integrons"):      
            full_path = os.path.join(root, file)
            file_list.append(full_path)

integron_region_df=pd.DataFrame(columns=['plasmid_id','start','end'])
for i in range(0,len(file_list)):
    integron_file=open(file_list[i])
    info=integron_file.readlines()
    integron_file.close()
    if "# No Integron found" in info[2]:
        continue
    else:
        integron_df=pd.read_csv(file_list[i],skiprows=2,sep='\t')
        plasmid_id=integron_df.loc[0,'ID_replicon']
        plasmid_df=pd.DataFrame(columns=integron_region_df.columns)
        for value, group in integron_df.groupby("ID_integron"):
            int_start=group[['pos_beg','pos_end']].min().min()
            int_end=group[['pos_beg','pos_end']].max().max()
            int_size=int_end-int_start
            temp_df=pd.DataFrame([{'plasmid_id':plasmid_id,'start':int_start,'end':int_end,'size':int_size}])
            plasmid_df=pd.concat([plasmid_df,temp_df],ignore_index=True)
        integron_region_df=pd.concat([integron_region_df,plasmid_df],ignore_index=True)

integron_region_df.to_csv(FILE_PATH,index=None)
```
