## Supplementary for "Synergistic interactions of mobile genetic elements shape the stepwise evolution of multidrug-resistant plasmids": Code_2_ARG_integron_in_or_out.py

```
import pandas as pd

ARG_loc=pd.read_csv(FILE_PATH)

integron_regions=pd.read_csv(FILE_PATH)

int_plasmid_list=sorted(set(integron_regions['plasmid_id']))
int_plasmid_ARG=ARG_loc[ARG_loc['Accession'].isin(int_plasmid_list)]
int_plasmid_ARG=int_plasmid_ARG.reset_index(drop=True)

for i in range(0,len(int_plasmid_ARG)):
    plasmid_id=int_plasmid_ARG.loc[i,'Accession'].strip()
    integron_for_plasmid=integron_regions[integron_regions['plasmid_id'].str.contains(plasmid_id, case=False, regex=False)]
    integron_for_plasmid=integron_for_plasmid.reset_index(drop=True)
    if len(integron_for_plasmid)==0:
        print('!!!')
    if "+" in int_plasmid_ARG.loc[i,'Location']:
        ARG_start=int(int_plasmid_ARG.loc[i,'Location'].split('+')[0].split('_')[2])
        ARG_end=ARG_start+int(int_plasmid_ARG.loc[i,'Location'].split('+')[1])-1
    elif "-" in int_plasmid_ARG.loc[i,'Location']:
        ARG_end=int(int_plasmid_ARG.loc[i,'Location'].split('-')[0].split('_')[2])
        ARG_start=ARG_end-int(int_plasmid_ARG.loc[i,'Location'].split('-')[1])+1
    int_plasmid_ARG.loc[i,'ARG_start']=ARG_start
    int_plasmid_ARG.loc[i,'ARG_end']=ARG_end
    overlap=False
    for j in range(0,len(integron_for_plasmid)):
        integron_start=int(integron_for_plasmid.loc[j,'start'])
        integron_end=int(integron_for_plasmid.loc[j,'end'])
        if integron_start<=ARG_start<=integron_end:
            overlap=True
            break
        elif integron_start<=ARG_end<=integron_end:
            overlap=True
            break
    int_plasmid_ARG.loc[i,'Integron-carrying']=overlap
    #print(overlap,integron_start,integron_end,ARG_start,ARG_end)

plasmid_ARG_in_out_count=pd.DataFrame()
for value, group in int_plasmid_ARG.groupby("Accession"):
    plasmid=group.iloc[0]['Accession']
    true_number=group['Integron-carrying'].sum()
    false_number=len(group)-true_number
    perc_in=true_number/len(group)
    perc_out=false_number/len(group)
    temp_df=pd.DataFrame([{'plasmid_id':plasmid,'in_integron':true_number,'out_integron':false_number,'perc_in':perc_in,'perc_out':perc_out}])
    plasmid_ARG_in_out_count=pd.concat([plasmid_ARG_in_out_count,temp_df],ignore_index=True)
    
plasmid_ARG_in_out_count.to_csv(FILE_PATH)
int_plasmid_ARG.to_csv(FILE_PATH)    
    
    
    
    
```
