## Supplementary for "Synergistic interactions of mobile genetic elements shape the stepwise evolution of multidrug-resistant plasmids": Code_3_ARG_unique_vs_total.py

```
import pandas as pd

input_df=pd.read_csv(FILE_PATH)

out_df=pd.DataFrame()
for name, group in input_df.groupby('Accession'):
    total=list(group['Element symbol'])
    unique=sorted(set(total))
    temp_df=pd.DataFrame(columns=['Accession','# of unique ARGs','# of total ARGs','unique/total'],data=[[name,len(unique),len(total),len(unique)/len(total)]])
    out_df=pd.concat([out_df,temp_df],ignore_index=True)

out_df.to_csv(FILE_PATH,index=None)
```
