## Supplementary for "Synergistic interactions of mobile genetic elements shape the stepwise evolution of multidrug-resistant plasmids": Code_4_OLS_RLM.py

```
import statsmodels.api as sm
import statsmodels.formula.api as smf
import pandas as pd

#Integron+ vs integron- slope and intercept
df=pd.read_csv(FILE_PATH)

model = smf.ols(formula="ARGs ~ IS26 * C(Integron)", data=df).fit()
print(model.summary())

model2 = smf.rlm('ARGs ~ IS26 * C(Integron)', data=df, M=sm.robust.norms.HuberT())
results = model2.fit()
print(results.summary())


#ISCR1 vs ISCR2 slope and intercept
df_2=pd.read_csv(FILE_PATH)

model3 = smf.rlm('ARGs ~ IS26 * C(ISCR)', data=df_2, M=sm.robust.norms.HuberT())
results_2 = model3.fit()
print(results_2.summary())
```
