## Supplementary for "Synergistic interactions of mobile genetic elements shape the stepwise evolution of multidrug-resistant plasmids": Supplementary_Information.pdf

### Inventory Summary

**Extended Data Fig. 1** Kolmogorov-Smirnov Test showed that the number of ARGs on integron-positive (n = 419) plasmids were significantly higher than those on integron-negative (n = 683) plasmids ($p < 0.0001$).

**Extended Data Fig. 2** Kolmogorov-Smirnov Test showed that the number of ARGs outside of the integron-associated structures (n = 419) are significantly higher than the number of total ARGs on integron-negative plasmids (n = 683) ($p < 0.0001$), even with the highly conservative assumption that all integrons contain two ARGs *sul1* and *qacEdelta1* in the 3'CS regions that are excluded from the integron regions. The asterisks mean the median values of the number of ARGs in each group.

**Extended Data Fig. 3** The gene annotation highlighting the ARGs, intI1, and the predicted integron regions for the 11 integron-positive plasmids collected from human and pig wastewater.

**Extended Data Fig. 4** The number of unique ARGs versus the number of total ARGs in the 1,102 ARG-carrying plasmids investigated in this study. The reference dash line means the number of unique ARGs equals the number of total ARGs. If a dot falls on the reference dash line, it means there are no replicated ARGs on the plasmid.

**Extended Data Fig. 5** The core genome tree, ARG pattern, and genome alignment of a plasmid lineage containing 10 plasmids. a) The core genome tree and number of different classes of ARGs in the 10 plasmids. Red solid circles indicate integron-positive plasmids, and red hollow circles indicate integron-negative plasmids. b) The genome alignment of the 10 plasmids. The IS26 elements are colored black in the alignment. Unaligned genes are colored grey.

**Extended Data Fig. 6** The numbers of complete integrons or In0 (found in 395 plasmids) were greater than the number of CALIN (found in 51 plasmids) on integron-positive plasmids, suggesting the integrons on plasmids were still able to capture new genes into their cassettes.

**Extended Data Fig. 7** The gene annotations of the IS26-flanked CALIN on four plasmids. a) NZ_CP041112.1; b) NZ_CP058835.1; c) NZ_CP095650.1; d) NZ_KX246266.1

**Extended Data Table 1.** The Chi-square test results for the presence and absence of integrons, IS26 elements, and ISCR elements for the 1,102 ARG-carrying plasmids (This table is provided in this file).

**Supplementary Table 1.** The information of the 1,102 ARG-carrying circular plasmids retrieved from PLSDB RefSeq (This table is provided as a separate Excel workbook).

**Supplementary Table 2.** Information of the 32 wastewater-derived Escherichia coli plasmids collected in this study (This table is provided as a separate Excel workbook).

**Supplementary Table 3.** All 3,211 circular plasmids retrieved from PLSDB RefSeq (This table is provided as a separate Excel workbook).

**Supplementary Table 4.** The antibiotic resistance gene (ARG) annotation results for the 1,102 circular ARG-carrying plasmids retrieved from PLSDB RefSeq (This table is provided as a separate Excel workbook).

**Supplementary Note 1.** Non-cassette ARGs on integron-positive plasmids were significantly more than ARGs on integron-negative plasmids (This note is provided in this file).

**Supplementary Note 2.** IS26-mediated rearrangements and IS26-associated homologous recombination events were observed across a lineage comprising 10 plasmids obtained from multiple samples (This note is provided in this file).

**Supplementary Software 1.** The customized Python codes used in this study (This software is provided as a separate zip file).

**Supplementary Data 1.** The output data for Ordinary Least Squares (OLS) and Robust Linear Model (RLM) regression analyses (This dataset is provided as a separate csv file).

**Supplementary Note 1. Non-cassette ARGs on integron-positive plasmids were significantly more than ARGs on integron-negative plasmids**

We asked if the increased number of ARGs in integron-positive plasmids solely were due to the cassette arrays that integrons carried. We classified the ARGs carried by each plasmid into two categories: inside integron (at least 1 bp overlap with the integron region) and outside integron (no sequence overlaps with the integron region). Because *sul1* and *qacEdelta1* were often considered as conserved structures on the 3' CS region of Class 1 integrons but the sequences of these two genes could not be counted into integron regions based on the algorithm of IntegronFinder 2.0, we further made a highly conservative assumption that all plasmids had two ARGs misclassified into the outside-integron group. We therefore increased the number of ARGs on integron-negative plasmids by two before conducting statistical tests to compare the number of ARGs outside integron regions on integron-positive plasmids and the number of ARGs carried by integron-negative plasmids. The statistical test results showed that the number of ARGs outside of integron regions on integron-positive plasmids were significantly higher than the number of ARGs carried by integron-negative plasmids with the two-gene-misclassification assumption, with medians of 6 and 4, respectively ($p < 0.0001$) (**Extended Data Fig. 2**). This finding suggests that the increased number of ARGs on integron-positive plasmids was not only due to the ARGs carried by integron cassette arrays, and there were additional strategies occurring.

**Supplementary Note 2. IS26-mediated rearrangements and IS26-associated homologous recombination events were observed across a lineage comprising 10 plasmids obtained from multiple samples.**

To investigate the distribution and organization of ARGs, integrons, and IS26 elements on plasmids, we analyzed 10 plasmids that shared highly similar backbone sequences with various numbers of ARGs and both integron presence and absence. The phylogenetic tree of the aligned core genes of the 10 plasmids with corresponding ARG counts is in **Extended Data Fig. 5a**. Varied ARG patterns were observed among the 10 plasmids. Trimethoprim resistance gene *dfrA14* which is often located in integron cassette arrays were found only on integron-positive plasmids as expected, as well as aminoglycoside resistance genes *aadA2*, *aac(6')-Ib*, and *rmtB1*.

The sequence alignment further illustrates the contribution of IS26 and integron in the ARG composition and rearrangement on these 10 plasmids (**Extended Data Fig. 5b**). Except for NZ_AP022370.1, the sulfonamide resistance gene *sul2* was flanked by two IS26 elements in inverted orientation in all nine other plasmids. In NZ_CP125348.1, NZ_CP088994.1, NZ_MN182748.1, and NZ_CP130664.1, the chloramphenicol resistance gene *catA2* was flanked by two IS26 elements in the same orientation and located immediately adjacent to the IS26-flanked gene cluster containing *sul2*, with the two regions sharing a common IS26 element. Similarly, in NZ_CP130664.1, NZ_CP107338.1, NZ_CP107331.1, NZ_CP102394.1, NZ_CP101783.1, and NZ_CP107293.1, the *dfrA14*-containing integron was flanked by two IS26 elements in the same orientation and located immediately adjacent to the same *sul2*-containing gene cluster, with a common IS26 element shared. In NZ_AP022370.1, the gene cluster flanked by two directly oriented IS26 elements was replaced by an intact integron carrying *aac(6')-Ib*, *bla*$_{IMP-6}$, and *aadA2* in its cassette array, with *sul1* and *qacEdelta1* in the 3' CS region. The *sul2*-containing gene cluster flanked by two inverted oriented IS26 elements was missing in NZ_AP022370.1. NZ_CP107293.1 had the highest ARG count (11 ARGs) among the 10 plasmids. According to the alignment figure, interestingly, NZ_CP107293.1 was a recombined plasmid containing both IncI and IncF replicons. The region in NZ_CP107293.1 that was not shared with the nine other plasmids was flanked by two directly oriented IS26 elements, suggesting the conserved sequences of the two IS26 elements were associated with the homologous recombination of the two plasmids. The additional ARGs on NZ_CP107293.1, such as $bla_{TEM-1}$, *fosA3*, $bla_{CTX-M-65}$, $bla_{KPC-2}$, and $bla_{SHV-12}$, were contributed by the recombined unshared regions on this plasmid. The diverse ARGs associated with integrons and IS26 elements in the same plasmid lineage suggest the important roles of both MGEs in ARG insertion and rearrangement on plasmids.

**Extended Data Table 1.** The Chi-square test results for the presence and absence of integrons, IS26 elements, and ISCR elements for the 1102 ARG-carrying plasmids ($p < 0.0001$ for all).

| | **ISCR +** | **ISCR -** | | **ISCR +** | **ISCR -** | | **Integron +** | **Integron -** |
|---|---|---|---|---|---|---|---|---|
| **IS26 +** | 194 | 446 | **Integron +** | 175 | 244 | **IS26 +** | 365 | 275 |
| **IS26 -** | 16 | 446 | **Integron -** | 35 | 648 | **IS26 -** | 54 | 408 |
| OR = 12.13<br>(95% CI: 7.16 - 20.53) | | | OR = 13.28<br>(95% CI: 8.98 - 19.64) | | | OR = 10.03<br>(95% CI: 7.25 - 13.87) | | |
