## Supplementary figures and images for "Synergistic interactions of mobile genetic elements shape the stepwise evolution of multidrug-resistant plasmids"

### Extended_data_figure_1_PLSDB_int_PN.tif

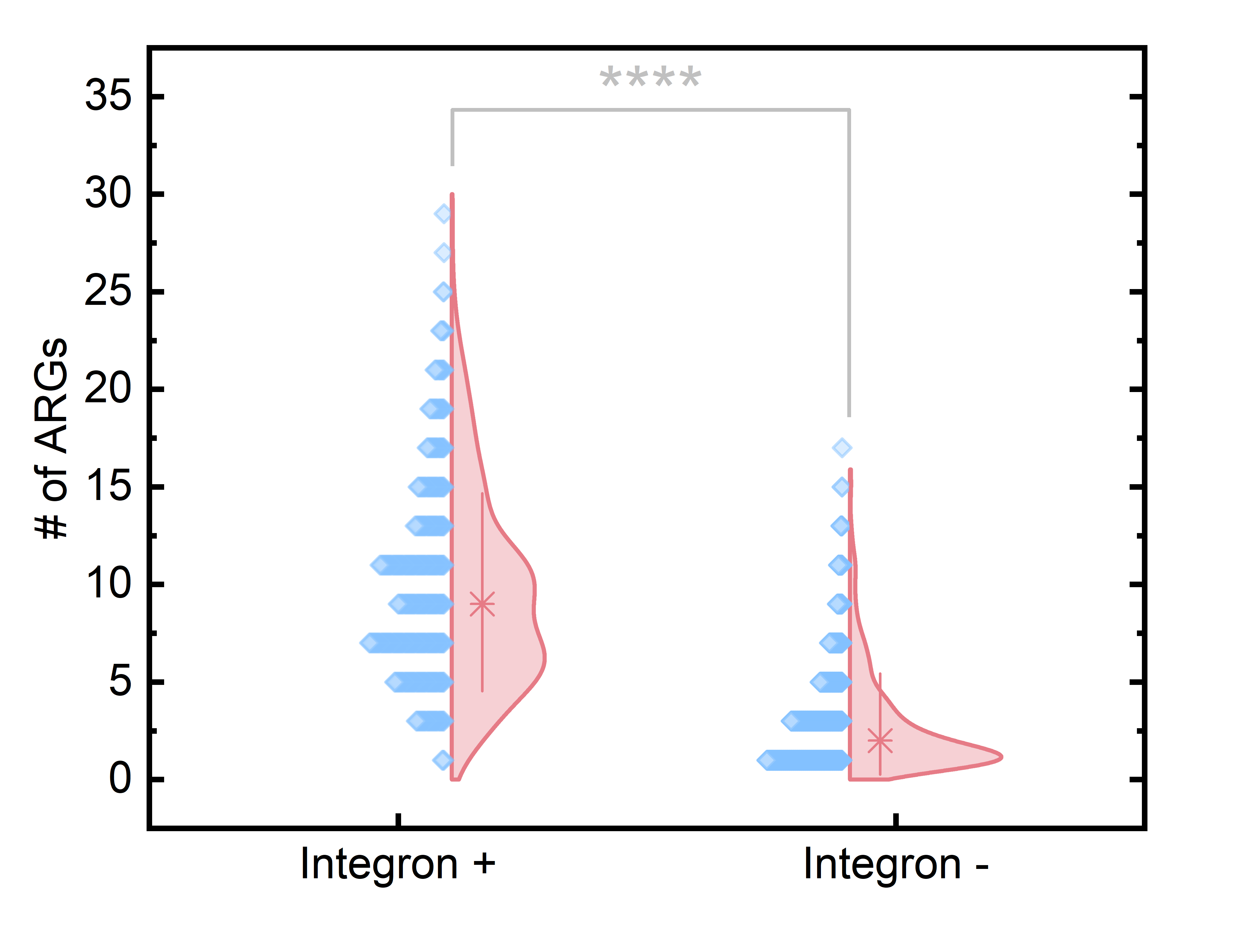

### Extended_data_figure_2_int_O_vs_N.tif

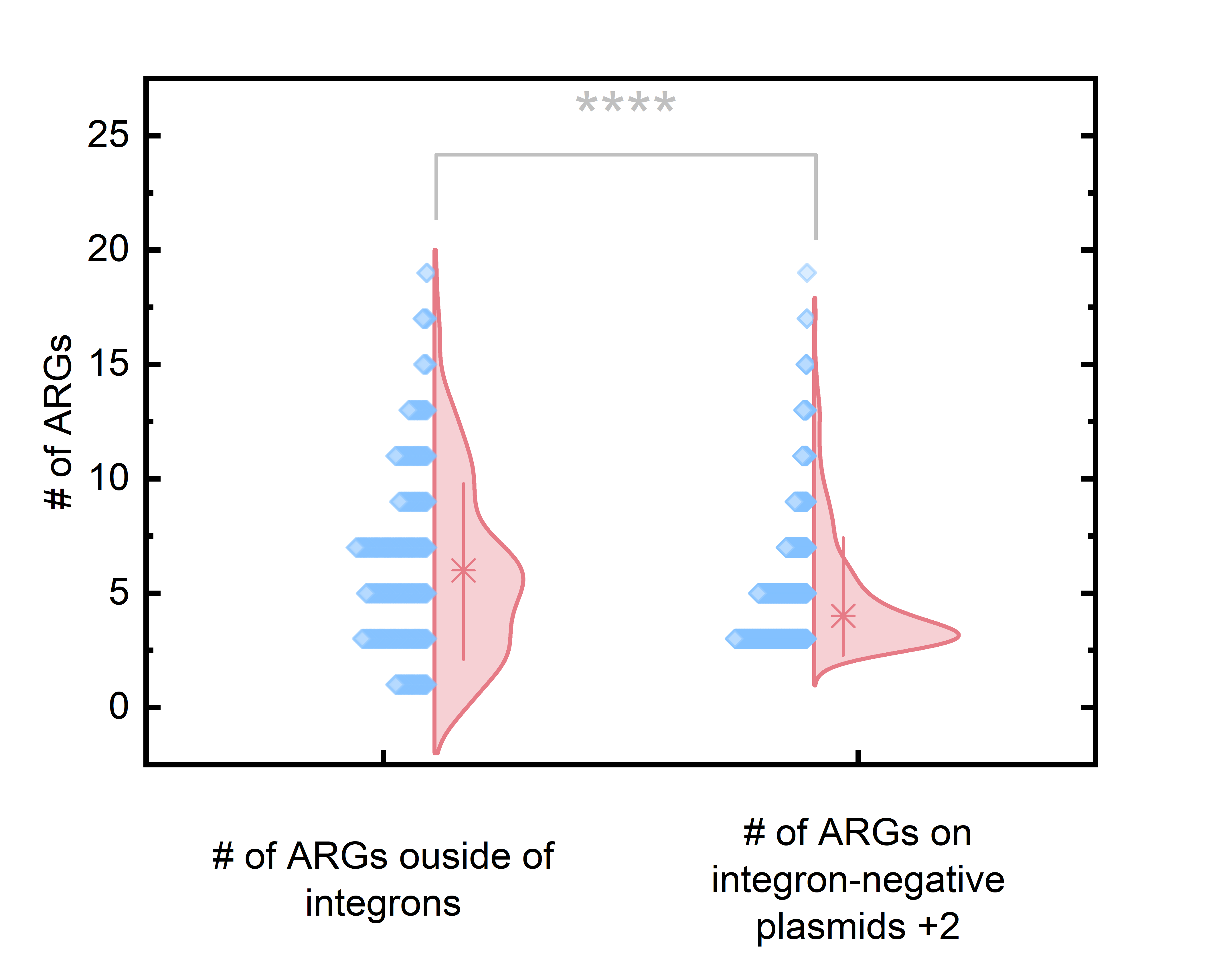

### Extended_data_figure_3_Ecoli_plasmid_annotation.tif

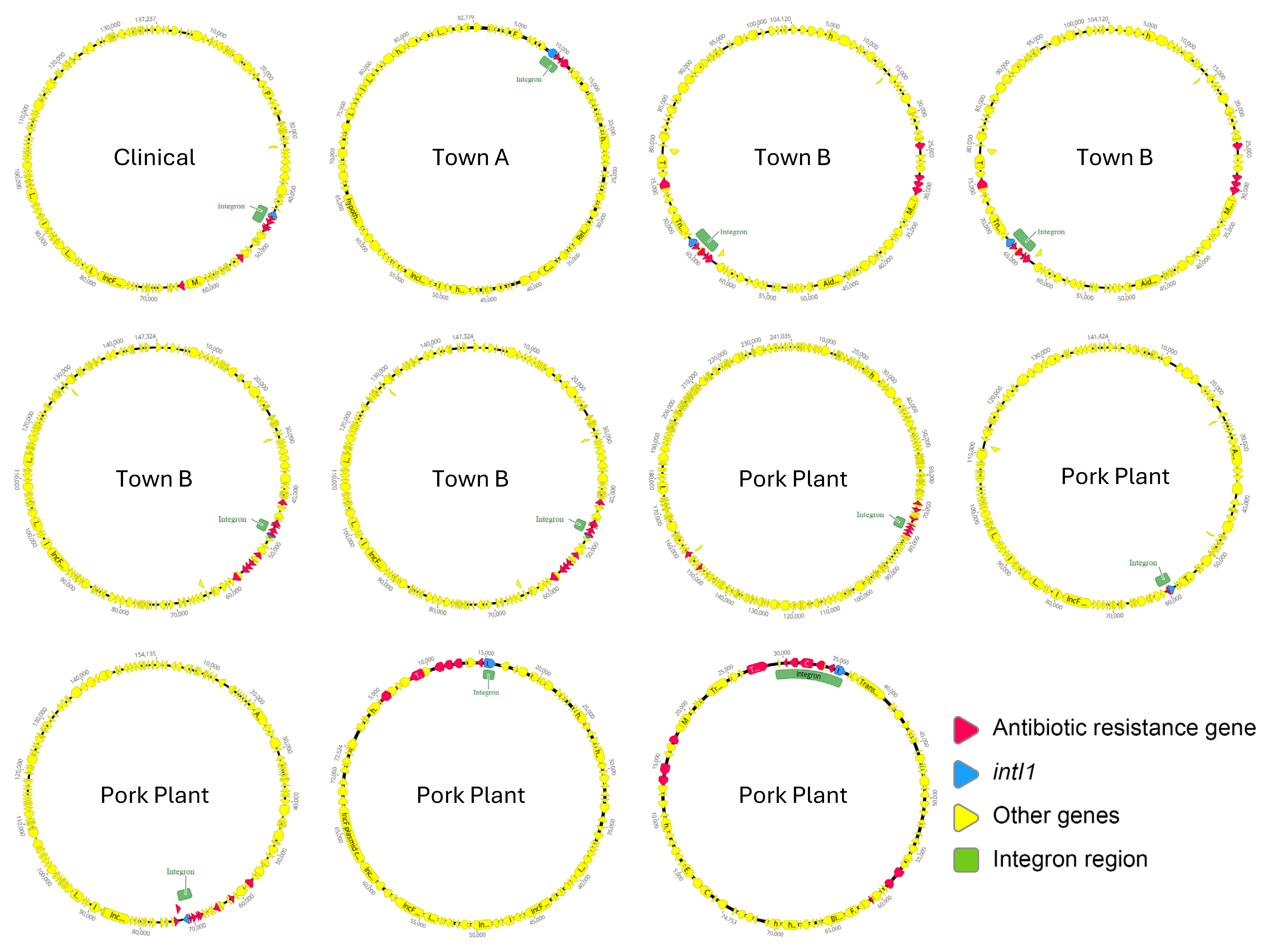

### Extended_data_figure_4_unique_vs_total.tif

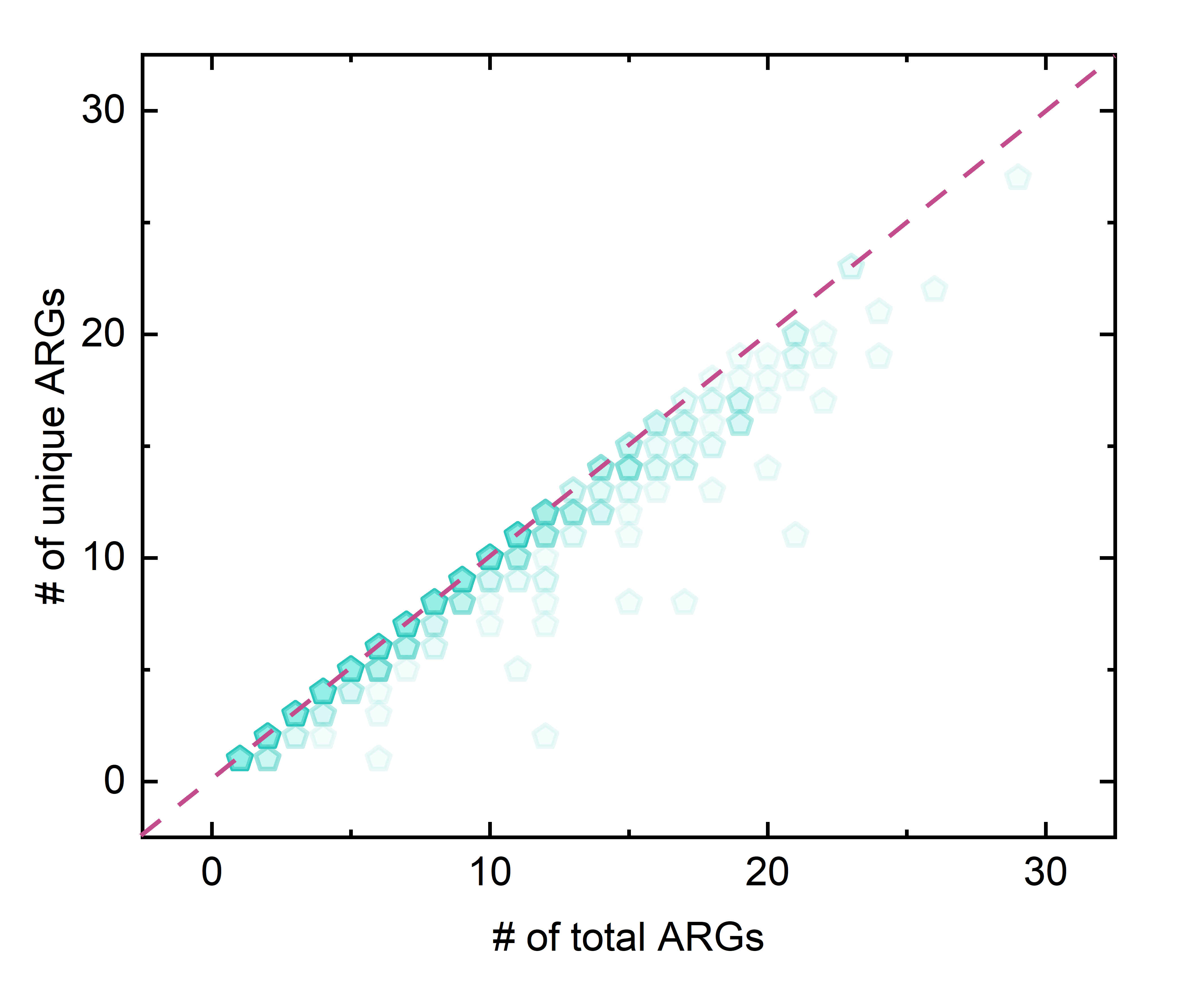

### Extended_data_figure_5_sub_cluster_tree_and_heatmap.tif

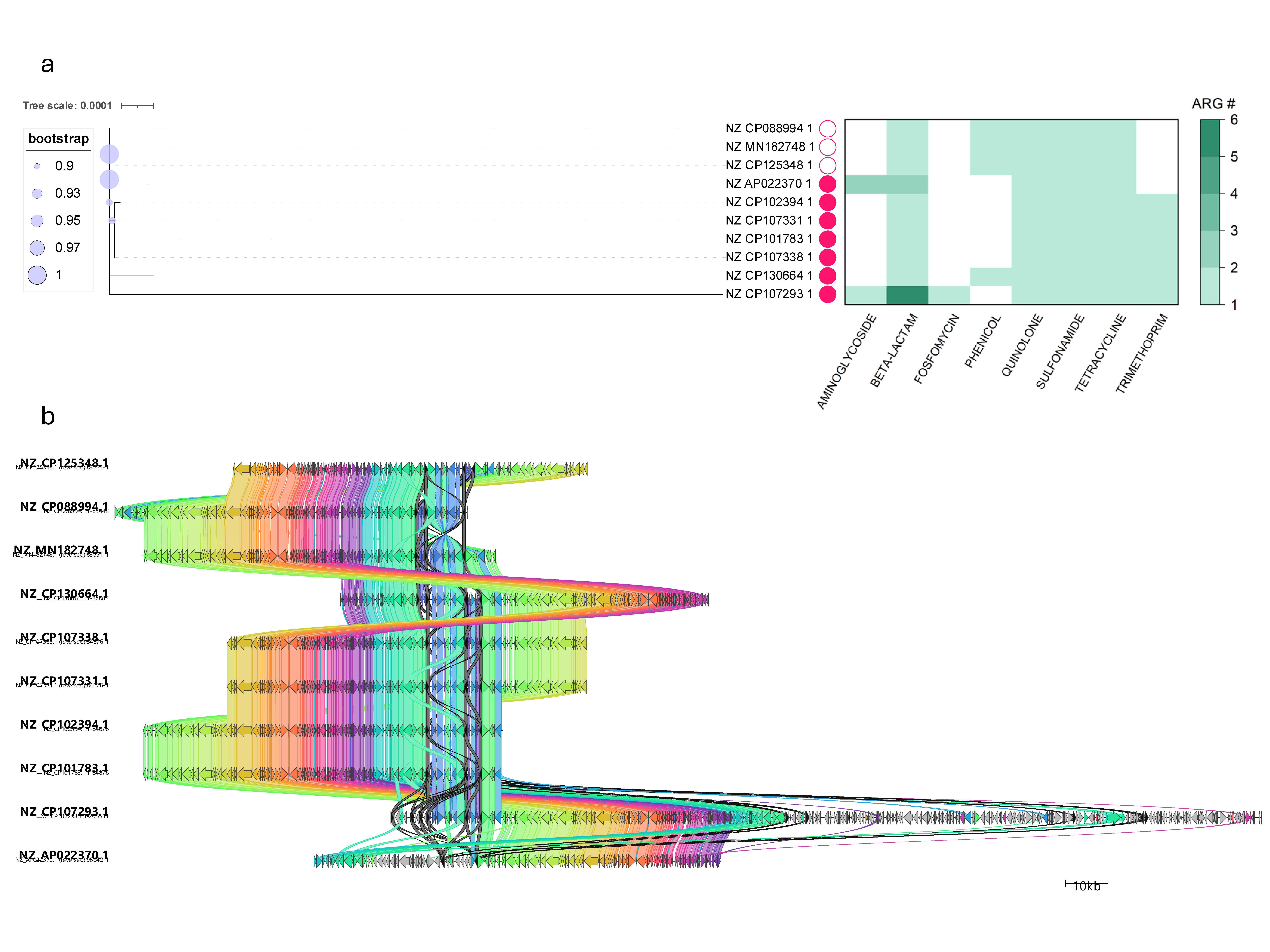

### Extended_data_figure_6_CALIN_In0.tif

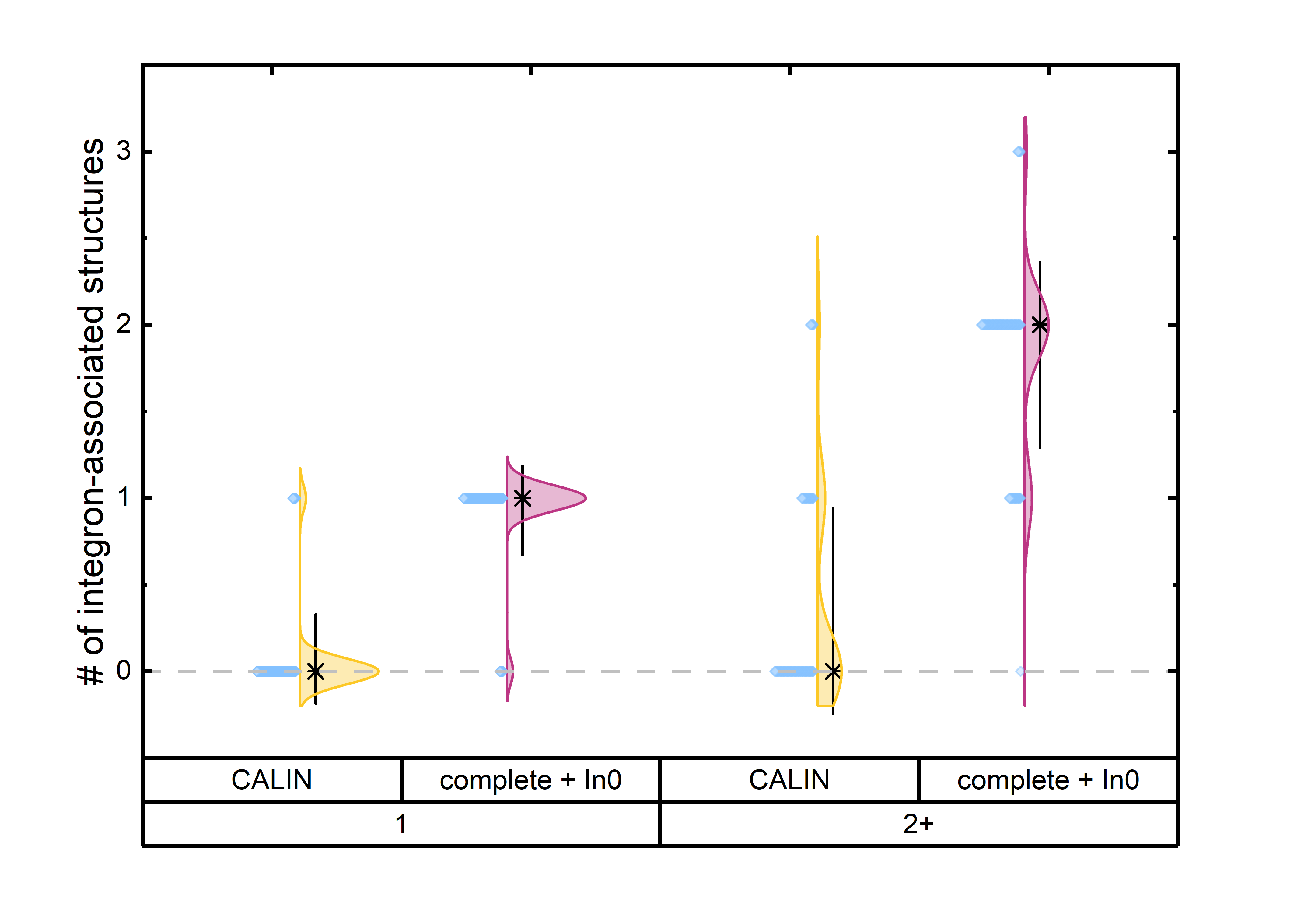

### Extended_data_figure_7_IS26_CALIN_annotation.tif

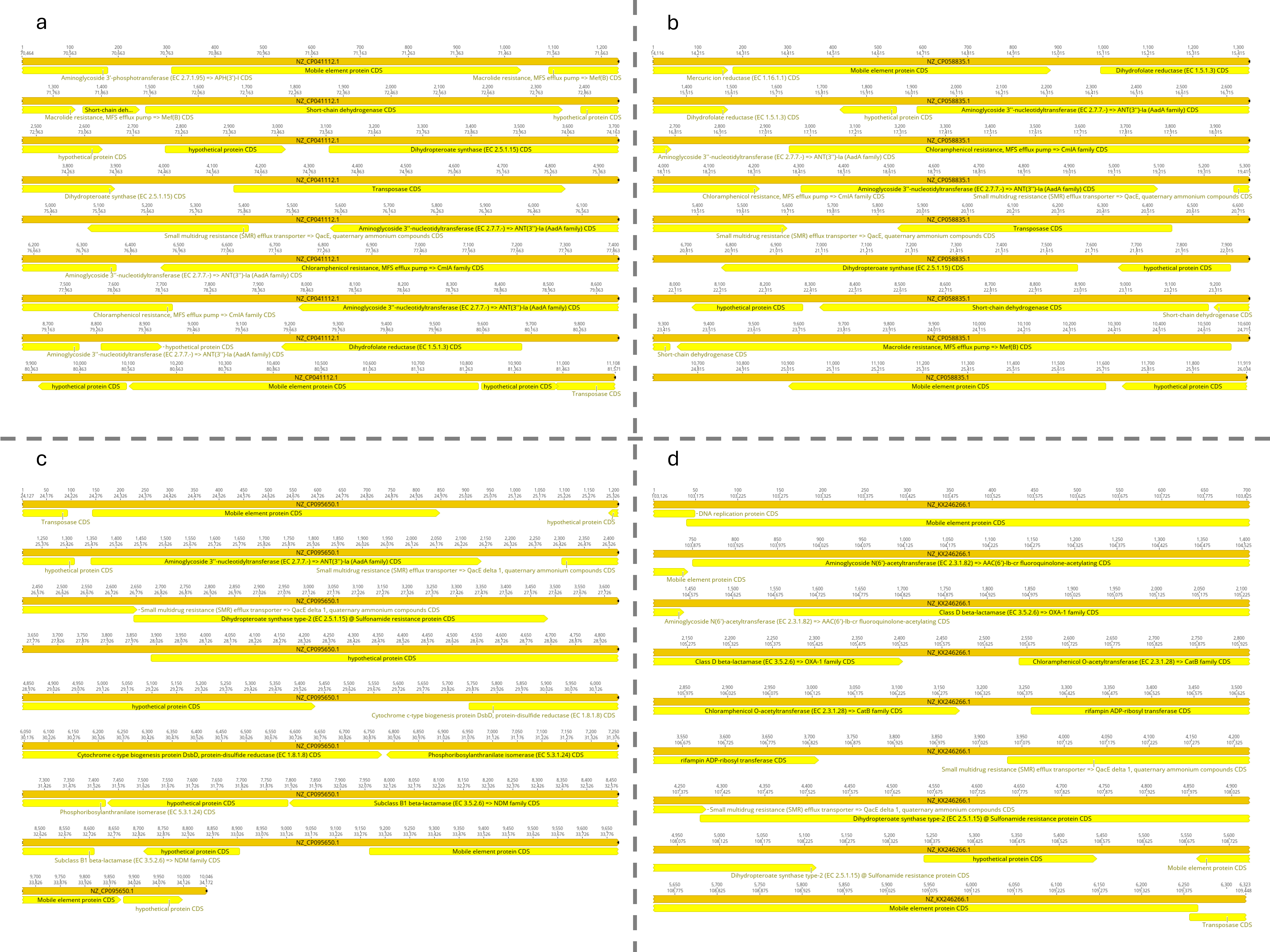
